# Cultural echoes in social bonding: Universal and culture-specific effects of movement synchrony and music on connectedness, likeability, and well-being

**DOI:** 10.64898/2026.08.31.748206

**Authors:** Mathias Klarlund, Hua Shan, Peter Vuust, Yi Du, Jan Stupacher

## Abstract

Social bonds are essential for well-being, and moving together to music is a universal means of promoting prosocial behavior and positive affect. However, the functions and interpretations of movement synchrony and music can vary across cultures and may depend on personal values. We examined how interpersonal synchrony and music affect social bonding in participants from individualistic (Denmark, N = 137) and collectivistic (China, N = 226) cultures. Participants watched videos of virtual stick figures - representing themselves (virtual-self) and an unknown person (virtual-other) - moving either synchronously or asynchronously to music or a metronome. After each video, participants rated three indicators of social bonding: their momentary feeling of social connectedness with the virtual-other (Inclusion of Other in the Self, IOS), likeability of the virtual-other, and well-being as the virtual-self. Personal values were measured using the Portrait Value Questionnaire. Music and synchrony promoted all three indicators of social bonding in both cultures, whereas cultural differences emerged in asynchronous conditions. Chinese participants rated the asynchronous virtual-other more likeable than Danish participants, aligning with the collectivistic emphasis on social harmony. Moreover, Chinese participants rated their own well-being higher than Danish participants when the virtual-self moved asynchronously, suggesting greater self-forgiveness in maintaining group harmony. The personal values of benevolence, conformity, and universalism interacted with movement synchrony and cultural background, underscoring the important role of values in interpretations of social interactions. Our findings highlight the universal prosocial effects of music and movement synchrony, while also emphasizing cultural nuances in how these effects are experienced.

## Introduction

Social connectedness is vital for well-being, influencing both psychological and physical health. Long-term social bonds contribute to emotional stability, stress resilience and overall life satisfaction (Umberson and Karas Montez, 2010), while social isolation is linked to lower survival rates (Holt-Lunstad et al., 2010). Social connectedness can also occur at shorter timescales, for example through momentary interpersonal synchrony, which can amplify perceptual and behavioral forms of social connection (Baek et al., 2025). Among various facilitators of social connectedness, music stands out as uniquely influential, being present across all known cultures (Trehub et al., 2015). Music fosters well-being by promoting human connection, shared experiences, and meaningful interactions (Schulkin & Raglan, 2014). It is therefore not surprising that interpersonal interactions featuring music enhance momentary, state-like social connectedness, as reflected in increased self-other integration, likability of others, and overall well-being (Stupacher et al., 2017). Over time, these short-term interactions may further contribute to more enduring social bonds, consistent with the suggestion that human musicality evolved to enhance synchronous, perceptual, and behavioral aspects of social connectedness (Loersch & Arbuckle, 2013; Savage et al., 2021).

In an increasingly interconnected world, understanding how cultural differences influence social bonding is essential. One fundamental framework for examining cultural differences is the individualism-collectivism dichotomy, which shapes social behavior and norms. Collectivistic cultures, such as China, prioritize community, social harmony and interdependence, with self-identity closely tied to social roles and relationships within the group (Han & Humphreys, 2016; Kitayama et al., 2007; Markus & Kitayama, 1991). In contrast, individualistic cultures, such as the United States or Denmark, emphasize personal identity, independence, individual achievement and uniqueness (Hofstede et al., 2010). These cultural orientations affect prosocial behaviors, including volunteering and helping others. For instance, Riemer et al. (2014) found that participants from collectivistic cultures demonstrate a higher propensity for group-oriented prosocial behaviors motivated by social obligations and a desire for harmony, while participants from individualistic cultures are more driven by personal preferences and individual goals. Similarly, Kuwabara et al. (2007) found that trust and cooperation in collectivistic cultures are rooted in group identity, whereas individualistic cultures rely more on individual assessments.

Despite music’s recognized role in fostering prosocial behavior and social connections, most research has focused on individualistic cultures (Cirelli, 2018; Clark and Giacomantonio, 2013; Stupacher et al., 2017; Wiltermuth & Heath, 2009). The extent to which cultural norms and personal values influence the prosocial effects of music-mediated interactions between individualistic and collectivistic cultures remains largely unexplored. While the individualism-collectivism framework provides valuable insights, it is often criticized for oversimplifying cultural complexity. Culture is a multifaceted and dynamic construct, encompassing diverse beliefs, values and practices beyond the binary distinction of individualism and collectivism (Triandis, 1995). Schwartz’s theory of basic values (Schwartz, 2012; Schwartz et al., 2012) provides a more nuanced approach, identifying values like benevolence, tradition, hedonism, and stimulation that conceptually align with individualistic and collectivistic cultural orientations (Schwartz, 1992; Schwartz et al., 2001; Schwartz & Cieciuch, 2022). For example, tradition and conformity are more salient in collectivistic cultures, emphasizing social harmony and conformity to social norms, whereas hedonism, which prioritizes personal pleasure and self-gratification, is more prominent in individualistic cultures (Schwartz, 1999). Schwartz’s values can be summarized into two overarching values: social focus and personal focus. Social focus values, comprising universalism, benevolence, tradition, conformity and security, align more closely with collectivist cultures, while personal focus values, comprising self-direction, stimulation, pleasure, achievement and power, are more closely associated with individualistic cultures (Schwartz, 1994; Schwartz, 1990). Importantly, individuals within any cultural background may embody values from both orientations (Oyserman et al., 2002; Schwartz, 1994). Although cultural background encompasses a wide range of influences, in cross-cultural research it is often operationalized through participants’ nationality, serving as a proxy for cultural dimensionality, such as individualism and collectivism (Hofstede et al., 2010).

This study investigates how cultural orientations (individualistic vs. collectivistic) and personal values, measured by the revised Portrait Value Questionnaire (PVQ-RR) (Schwartz et al., 2016; Schwartz & Butenko, 2014; Schwartz et al., 2012), affect different indicators of social bonding in Danish and Chinese participants. Using a video paradigm, participants watched stick figures representing themselves (virtual-self) and another individual (virtual-other) moving either synchronously or asynchronously to music or a metronome (Stupacher et al., 2017; 2020; 2022). Participants then rated three indicators of social bonding in our experimental design: momentary feelings of social connectedness (Inclusion of Other in the Self; Aron et al., 1992), likeability of the virtual-other, and well-being of the virtual-self. We hypothesize that, consistent with prior research in Western samples (Austria and Germany) (Stupacher et al., 2017), movement synchrony and music will increase social connectedness, likeability, and well-being in both cultural contexts. Additionally, we expect that cultural orientation will shape responses to asynchronous movement in two ways. First, Chinese participants, given the cultural emphasis on social harmony, will show less negative evaluation of asynchronous others than Danish participants, reflecting a culturally guided tendency to preserve social cohesion. Second, Chinese participants will rate their own well-being lower when the virtual-self moves asynchronously, reflecting heightened self-criticism for disrupting harmony.

## Method

### Participants

Data were collected online from 413 participants: 250 Chinese and 163 Danish. We excluded 41 participants who failed the attention check and 9 Danish participants over the age of 40 to ensure comparable age distributions between the groups. The final sample included 363 participants: 226 Chinese (116 female, 110 male, M = 20.3 years, SD = 2.1) and 137 Danish (90 female, 45 male, 2 other, M = 25.0 years, SD = 3.3). Throughout this study, nationality refers to participants’ country of birth and upbringing—either Denmark or China—which serves as a proxy for their cultural background, the conceptual focus of this study. Importantly, all participants were residing in their country of birth at the time of the study (China or Denmark). Based on previous studies using similar paradigms (Stupacher et al., 2017; 2020; 2022), we aimed to recruit a minimum of 150 participants per nationality before data exclusion to ensure a sufficiently large final sample after applying exclusion criteria. All participants provided digital informed consent, and the study was approved by the Institutional Review Board at the … (…).

### Stimuli and Design

Participants watched and rated six 12-second videos displaying two walking stick figures, a black figure representing the participant themselves (virtual-self) and a blue figure representing an unknown person (virtual-other). Similar stimuli have been used in previous studies (Stupacher et al., 2017, 2020, 2022) and participants are generally able to identify with the stick figures (Stupacher et al., 2017). Although the stick-figure design reduced external validity, it minimized biases related to physical attributes such as gender, age, or ethnicity, thereby increasing internal validity. We manipulated two independent variables in these videos: movement synchrony and audio. Movement synchrony had three levels: (1) both figures synchronized with the beat, (2) virtual-self synchronized and virtual-other asynchronized (self sync / other async), and (3) virtual-self asynchronized and virtual-other synchronized (self async / other sync) (Figure 1A-C). Audio had two levels: music and metronome (Figure 1D-E). Including both music and metronome allowed us to disentangle effects of rhythmic structure from those specifically attributable to the socio-emotional and affective qualities of music. The music was a generic instrumental piece composed for a previous study (Stupacher et al., 2016) and therefore unfamiliar to the participants. Both music and metronome had a tempo of 94.3 beats per minute (BPM; 636 ms per beat). Synchronized figures stepped at 94.3 bpm, hitting the ground on the beats of the music or the metronome, with a “dust cloud” on the ground visually marking each step on the beat (Figure 1). Asynchronized figures stepped at 80 bpm (750 ms per beat) with a 125 ms delayed first step, creating a constantly changing phase relationship between the synchronized and asynchronized steps. The videos are available online at https://researchbox.org/3886&PEER_REVIEW_passcode=LVBGRI.

**Figure 1:**
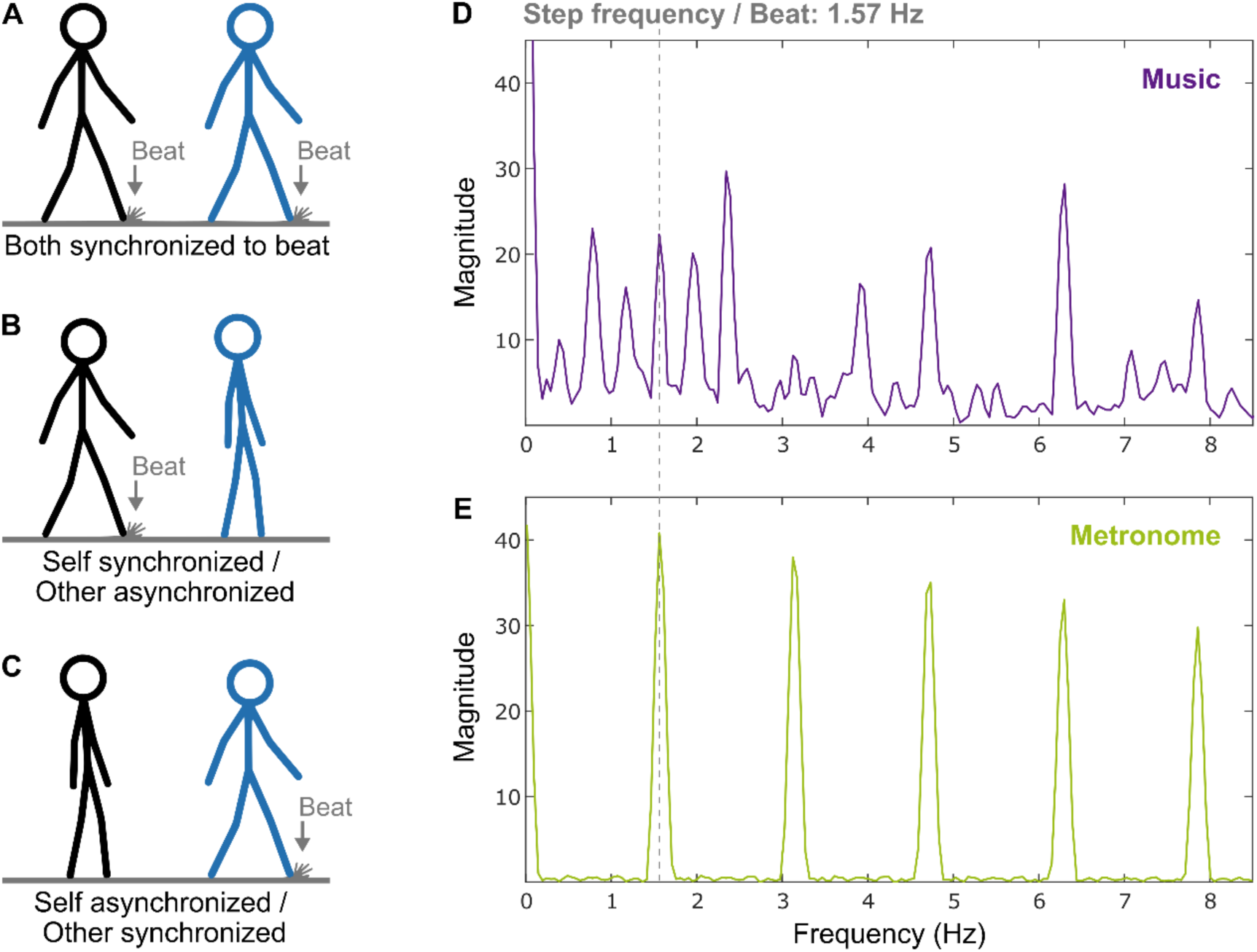
Experimental design and information of stimuli. Three levels of the movement manipulation: Both figures synchronized with the beat (A), virtual-self synchronized / virtual-other asynchronized (B), and virtual-self asynchronized / virtual-other synchronized (C). Frequency spectra of the event-onset interval series of the music (D) and the metronome (E) computed with the MIR Toolbox (Lartillot & Toiviainen, 2007) for MATLAB (Mathworks, Natick, MA).

### Procedure and Ratings

Stimuli were presented and data were collected online on soscisurvey.de (SoSci Survey, Munich, Germany). Participants watched the 6 videos twice: once in a first block presented in random order, and once again in a second block with a new random order. We used two blocks to reduce within-participant variability and increased the stability of the estimates. After each video, participants rated three variables: social connectedness between the virtual-self and virtual-other using the Inclusion of Other in the Self (IOS) scale (Aron et al., 1992), the likeability of the virtual-other (“How much did you like the other (blue) figure?” ranging from “not at all” to “very much”), and their well-being as the virtual-self (“How did you feel in this situation as the black figure?” ranging from “not good at all” to “very good”). All ratings were given on a continuous slider ranging from 0 on the left to 100 on the right, with participants unable to see the numerical values. To ensure attentiveness of participants, one additional video served as an attention check, where a voice instructed participants to move all three rating sliders to the maximum value (100), instead of playing music or metronome sounds. Participants who failed to follow this instruction (i.e., did not rate at least 95 on all three scales) were excluded from analyses: 9.5% of Chinese and 9.4% of Danish participants.

After the video rating, participants rated how much they liked the music and how familiar they were with the type of music. Additionally, participants filled out several questionnaires: the brief form of the Interpersonal Reactivity Index (B-IRI) (Ingoglia et al., 2016), with empathy operationalized as the overall score of the B-IRI, the revised Portrait Value Questionnaire (PVQ-RR) (Schwartz et al., 2012; Schwartz & Cieciuch, 2022), the sensorimotor and social reward subscales of the Barcelona Music Reward Questionnaire (BMRQ) (Mas-Herrero et al., 2013), and the musical training subscale of the Goldsmiths Musical Sophistication Index (Gold-MSI) (Müllensiefen et al., 2013, 2014).

For Danish participants, instructions, ratings, and questionnaires were in English, except for the PVQ-RR, which was translated into Danish by author xxx in consultation with a professional Danish-English translator and the PVQ-RR’s original author (Schwartz et al., 2012; Schwartz and Cieciuch, 2022). Instructions and ratings for Chinese participants were translated into Mandarin Chinese by authors xxx and xxx, and all questionnaires used were validated versions in simplified Chinese. Specifically, the IRI was adapted from Zhang et al. (2010), the PVQ-RR was translated by Chinese researchers in collaboration with Schwartz, and its reliability and validity were established through confirmatory factor analysis (CFA) and multidimensional scaling (MDS) analyses (Schwartz & Cieciuch, 2022), the BMRQ was translated and validated by Jiang et al. (2025), and a simplified Chinese version of the Gold-MSI was validated by Li et al. (2024).

### Statistical Analysis

#### Effects of Nationality

To test the effect of *movement synchrony*, *audio* (music vs. metronome), and *nationality* (Chinese vs. Danish) on the ratings of social connectedness (IOS), likeability of the virtual-other, and well-being as the virtual-self, we formulated three linear mixed-effects models, one for each rating variable, using the lme4 package (Bates et al., 2015) in R/RStudio (https://www.r-project.org; https://posit.co/):

*rating variable ∼ synchrony * audio * nationality + empathy + bmrqSR + (synchrony * audio|participant ID) + (1|block)*

The fixed effects of *empathy* and *BMRQ social reward* were included as control variables in the models as they are important for social interactions and significantly differed between the two cultural groups (Figure 2B, 2C). The models also included random slopes and intercepts for *movement synchrony* and *audio*, as well as random intercepts for participants and blocks. To assess the significance of the fixed effects, we performed type III analysis of variance with Satterthwaite’s approximation for estimating the degrees of freedom. Pairwise comparisons were computed with the *emmeans* package for R (Lenth et al., 2020), with Tukey’s method for multiple comparison adjustments.

**Figure 2:**
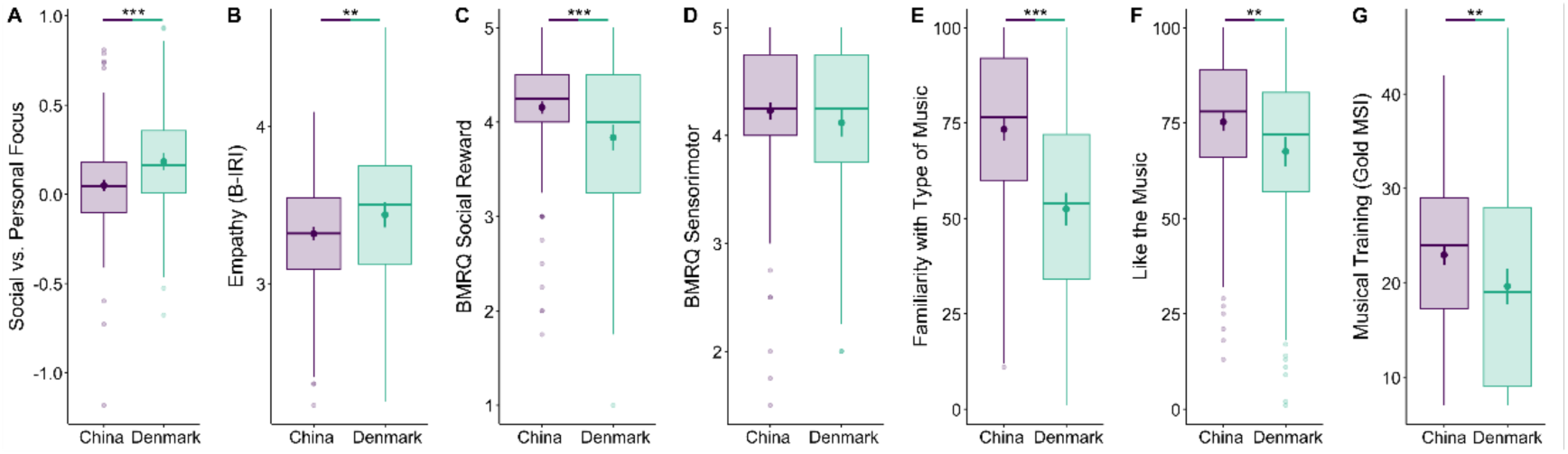
Control variables of participants values, traits, music ratings, and musical training. Nationality groups were compared with Welch two-sample t-tests. *** *p* < .001, ** *p* < .01.

#### Effects of Personal Values (PVQ-RR)

To address the individual variability often overlooked by the individualism-collectivism dichotomy, we conducted two additional analyses. First, we tested whether replacing nationality in the linear mixed-effect models with the two broad value categories from the PVQ-RR, social focus and personal focus, resulted in better model fits. Second, we examined in an exploratory analysis which of the 10 basic values in the PVQ-RR were most relevant for the ratings of social connectedness (IOS), likeability of the virtual-other, and well-being as the virtual-self. We chose the aggregate 10 basic values (rather than all the 19 PVQ-RR sub-dimensions) for simplicity and to maintain comparability ease with the literature.

For the first analysis, we compared the null model (*rating variable ∼ synchrony * audio + empathy + bmrqSR + (synchrony * audio|participant ID) + (1|block)*) with three models including either interactions between *synchrony * audio * nationality*, *synchrony * audio * social focus*, or *synchrony * audio * personal focus*.

For the second analysis, we fitted three linear mixed-effect models, one for each rating variable. Each model included all 10 basic values of the PVQ-RR and interactions of each basic value with *movement synchrony* and *nationality*. The models also included random intercepts and slopes for *movement synchrony*, and random intercepts for *audio*, *blocks*, and participants. We omitted the variable audio for this analysis, as it showed no significant interaction with nationality in the main analysis. The significance of the effects was assessed with type III analysis of variance with Satterthwaite’s approximation for estimating the degrees of freedom. Pairwise comparisons were computed with the *emmeans* package and the Tukey method for multiple comparison adjustments.

## Results

### Effects of Nationality

#### Control Variables

The analysis of participants’ values using the PVQ-RR showed that Chinese participants had a stronger social focus than Danish participants (*t*(257.6) = -4.44, *p* < .001, *d* = 0.49; Figure 2A), confirming our hypothesis about cultural values. This comparison was based on adjusted PVQ-RR values correcting for individual differences in the use of the response scale, as suggested by Schwartz (2016). Additional analyses revealed that Danish participants scored higher on empathy (estimated M = 3.44) than Chinese participants (estimated *M* = 3.32; *t*(217.2) = 2.61, *p* = .001, *d* = 0.29; Figure 2B), and that Chinese participants scored higher on social reward from music (estimated M = 4.15) than Danish participants (estimated *M* = 3.84; *t*(207.1) = 4.07, *p* < .001, *d* = 0.46; Figure 2C). The sensorimotor dimension of the BMRQ did not significantly differ between cultures (*t*(239.1) = 1.46, *p* = .144; Figure 2D). Chinese participants reported higher familiarity with and liking of the music used in the study (*t*(260.3) = 8.02, *p* < .001, *d* = 0.88 and *t*(241.0) = 4.07, *p* = .001, *d* = 0.37; Figures 2E and 2F) and were musically more trained than Danish participants (*t*(220.2) = 3.02, *p* = .003; Gold-MSI musical training scores of 22.9 and 19.6, respectively; Figure 2G). Importantly, adding familiarity with the music, liking of the music, or musical training as control variables to the models of IOS, likeability, and well-being ratings did not alter the significance or non-significance of any of the effects: *Movement Synchrony*, *Audio*, *Nationality*, *Nationality* × *Movement Synchrony*, *Nationality* × *Audio*, *Movement Synchrony* × *Audio*, and *Nationality* × *Movement Synchrony* × *Audio*. Correlations between musical training and the three dependent variables were positive but weak (musical training × IOS: *r*(361) = .13, *p* = .019; musical training × likeability of virtual other: *r*(361) = .11, *p* = .039; and musical training × well-being of virtual self: *r*(361) = .03, *p* = .574). Ratings of social connectedness, likeability of the other, and well-being as self did not differ between female and male participants (*t*(333.54) = -0.53, *p* = .595; *t*(308.61) = 1.57, *p* = .118; *t*(327.22) = 1.10, *p* = .272; respectively; Figure S1).

#### Social Connectedness

Social connectedness ratings showed significant main effects of movement synchrony, audio, and nationality, as well as an interaction between movement synchrony and nationality (Table 1). Social connectedness was rated higher in the synchronous condition than in both asynchronous conditions (both *p* < .001), with no significant difference between the two asynchronous conditions (*p* = .520). Overall, social connectedness was rated higher by Chinese participants compared to Danish participants (estimated mean difference (EMD) = 8.4, *p* < .001) and higher with music compared to metronome (EMD = 4.1, *p* < .001, Figure 3A). The interaction between movement synchrony and nationality indicated that in the two asynchronous conditions, social connectedness was rated higher by Chinese participants (EMDs 14.4 and 14.0), whereas no significant difference was observed in the synchronous condition (EMD = -3.4, Figure 4A). Higher empathy and social reward sensitivity from music were positively associated with higher ratings of social connectedness (Table 1, estimated marginal means of linear trends: 5.20 and 2.89, respectively). In summary, music enhanced social connectedness overall, replicating Stupacher et al. (2017), and Chinese participants reported feeling more connected to the asynchronously moving virtual partner than Danish participants. This pattern may reflect a greater tolerance for misalignment to preserve group cohesion.

**Figure 3:**
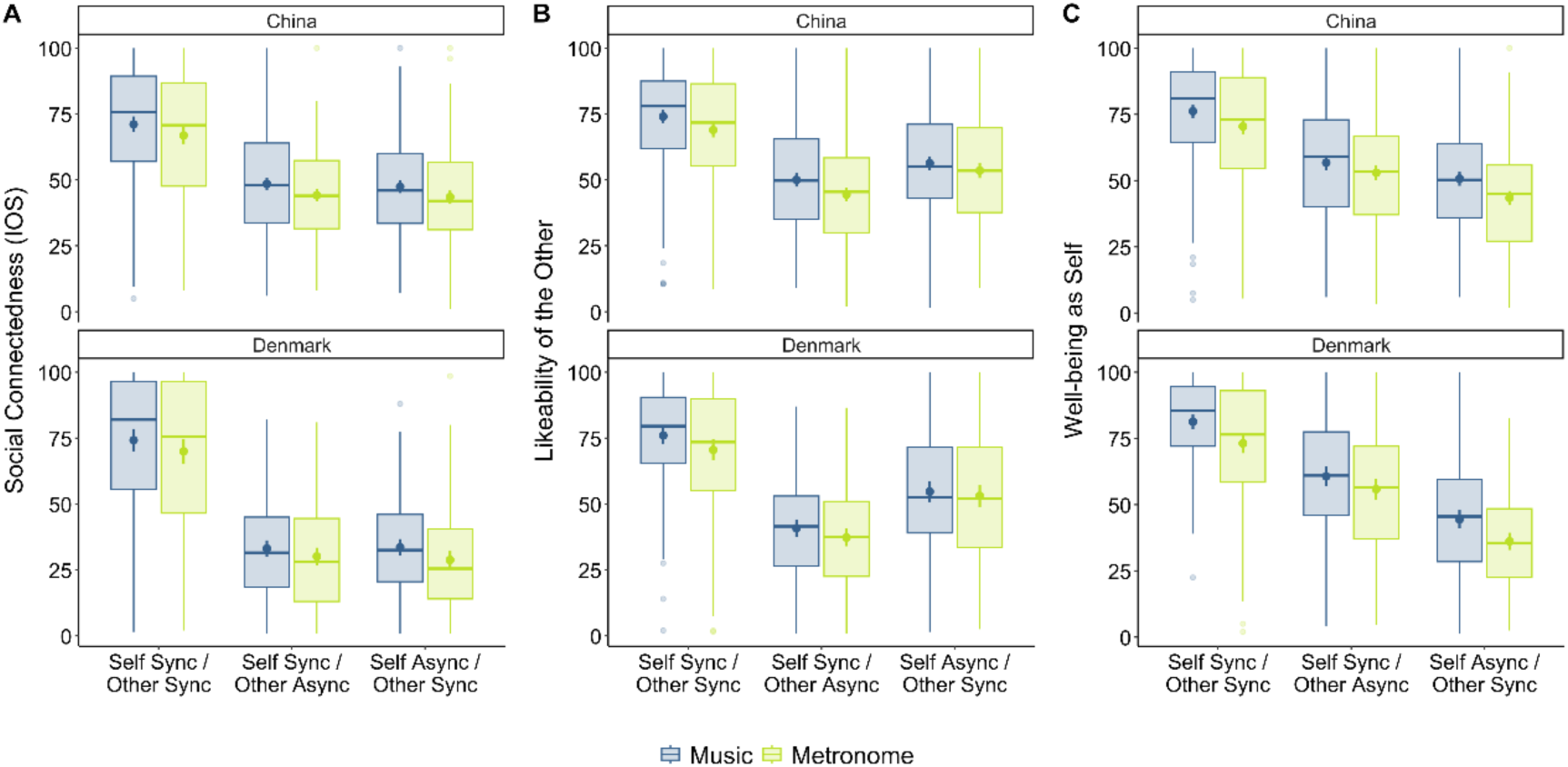
Cross-cultural difference of self-rating in different audio conditions. Mean ratings (●) with 95% confidence intervals (error bars) of social connectedness (A), likeability of the virtual other (B), and well-being as virtual self (C) for the three movement synchrony conditions and the two audio conditions (music in dark blue and metronome in bright green). Boxes of the boxplots display the lower quartile, median, and upper quartile. Whiskers of the boxplots display data within 1.5 times the inter-quartile-range below the lower quartile and above the upper quartile. Dots represent datapoints outside this range.

**Figure 4:**
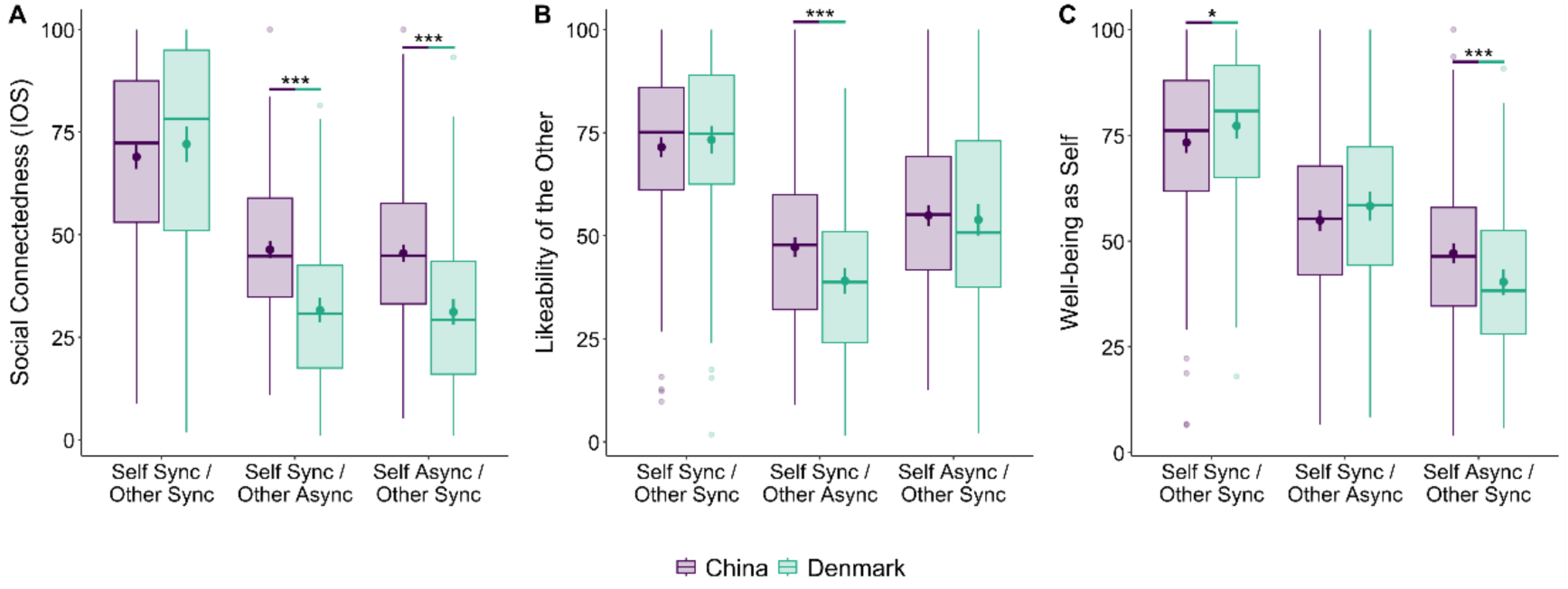
Cross-cultural difference of self-rating. Differences between Chinese and Danish participants in ratings of social connectedness (A), likeability of the virtual other (B), and well-being as virtual self (C) for the three movement synchrony conditions. Dots (●) represent means and error bars represent 95% confidence intervals. Boxes of the boxplots display the lower quartile, median, and upper quartile. Whiskers of the boxplots display data within 1.5 times the inter-quartile-range below the lower quartile and above the upper quartile. Dots represent datapoints outside this range. Data are averaged over the two audio conditions (music and metronome). *** *p* < .001 and * *p* < .05 in pairwise comparisons computed with *emmeans*.

**Table 1.** Results of the three individual linear mixed-effects models on social connectedness, likeability of the virtual other, and well-being as the virtual self.

| Effect | Social Connectedness (IOS) |  |  | Likeability of Virtual Other |  |  | Well-being as Virtual Self |  |  |
| --- | --- | --- | --- | --- | --- | --- | --- | --- | --- |
|  | *df* | *F* | *p* | *df* | *F* | *p* | *df* | *F* | *p* |
| Movement Synchrony | 2, 363.6 | 324.85 | < .001*** | 2, 361.0 | 301.91 | < .001*** | 2, 361.1 | 305.15 | < .001*** |
| Audio | 1, 361.8 | 47.33 | < .001*** | 1, 361.4 | 35.92 | < .001*** | 1, 362.1 | 72.63 | < .001*** |
| Nationality | 1, 361.2 | 23.27 | < .001*** | 1, 363.5 | 2.86 | .092 | 1, 363.1 | 0.05 | .822 |
| Nationality × Movement Synchrony | 2, 363.6 | 25.03 | < .001*** | 2, 361.0 | 7.84 | < .001*** | 2, 361.1 | 11.66 | < .001*** |
| Nationality × Audio | 1, 361.8 | 0.01 | .914 | 1, 361.4 | 0.53 | .465 | 1, 362.1 | 1.09 | .298 |
| Movement Synchrony × Audio | 2, 2192.4 | 0.26 | .773 | 2, 2575.3 | 4.09 | .017* | 2, 1589.1 | 4.27 | .014* |
| Nationality × Movement Synchrony × Audio | 2, 2192.4 | 0.57 | .568 | 2, 2575.3 | 0.58 | .561 | 2, 1589.1 | 0.23 | .793 |
| Empathy | 1,359.0 | 6.40 | .012* | 1,359.1 | 12.81 | < .001*** | 1,359.2 | 3.88 | .049* |
| BMRQ Social Reward | 1,359.0 | 5.48 | .020* | 1,359.1 | 1.94 | .165 | 1,359.2 | 3.02 | .083 |

Type III analysis of variance with Satterthwaite's method for estimating degrees of freedom. \*\*\* $p < .001$ ,
\* $p < .05$ .

#### Likeability of the Virtual Other

Likeability ratings of the virtual other showed significant main effects of movement synchrony and audio (Table 1). Likeability was rated higher in the synchronous condition compared to both asynchronous conditions (both *p* < .001), and higher in the self async / other sync condition than the self sync / other async condition (*p* < .001). Overall, likeability was rated higher with music than with the metronome (EMD = 4.0, *p* < .001, Figure 3B). A significant interaction between movement synchrony and nationality (Table 1) indicated that in the self sync / other async condition, likeability was rated higher by Chinese participants compared to Danish participants (EMD = 8.5, *p* < .001), whereas no significant differences were found between nationalities in the other two movement synchrony conditions (Figure 4B). Additionally, movement synchrony interacted with audio (Table 1, Figure 3B). Pairwise comparisons revealed that although likeability was rated significantly higher with music in all three movement synchrony conditions, the differences between music and metronome were higher in the self sync / other sync (EMD = 5.3) and self sync / other async (EMD = 4.5) conditions than in the self async / other sync condition (EMD = 2.2). Higher empathy was positively associated with higher likeability ratings (Table 1, estimated marginal mean of linear trend: 6.89). In sum, the likeability ratings mirrored the pattern of social connectedness ratings: music enhanced likeability overall, and Chinese participants evaluated the asynchronously moving virtual partner more favorably than Danish participants. This difference may indicate a stronger cultural tendency to prioritize group cohesion and accommodate interpersonal misalignment.

#### Well-being as the Virtual Self

Well-being ratings as the virtual self showed significant main effects of movement synchrony and audio (Table 1). Well-being was rated higher in the synchronous condition compared to the two asynchronous conditions (both *p* < .001), and higher in the self sync / other async condition compared to the self async / other sync condition (*p* < .001). Overall, well-being was rated higher with music compared to the metronome (EMD = 6.4, *p* < .001, Figure 3C). A significant interaction between movement synchrony and nationality (Table 1) indicated that in the self sync / other sync condition, well-being was rated higher by Danish compared to Chinese participants (EMD = -4.1, *p* = .042), while in the self async / other sync condition, well-being was rated higher by Chinese compared to Danish participants (EMD = 6.6, *p* = .001); the difference between the two nationalities in the self sync / other async condition was not significant (EMD = -3.6, *p* = .094; Figure 4C). Movement synchrony also interacted with audio (Table 1, Figure 3C). Pairwise comparisons revealed that although well-being was rated significantly higher with music in all three movement synchrony conditions, the differences between music and metronome were higher in the self sync / other sync (EMD = 7.0) and self async / other sync (EMD = 7.8) conditions than in the self sync / other async condition (EMD = 4.3). Higher empathy was positively associated with higher ratings of well-being (Table 1, estimated marginal mean of linear trend: 3.63). Therefore, well-being was generally rated higher with music compared to the metronome, and Chinese participants rated their own well-being as asynchronized figure higher than Danish participants. This pattern, consistent with the social connectedness and likeability results, may reflect a cultural tendency to prioritize group harmony over individual performance.

### Effects of Personal Values (PVQ-RR)

#### Nationality vs. Social and Personal Focus

For all three ratings, social connectedness, likeability of the virtual other, and well-being as the virtual self, adding an interaction term for nationality significantly improved the model fit compared to the null model (all *Chi*^2^ > 22 and all *p* ≤ .001), with reductions of the Akaike information criterion (AIC) by 67, 10, and 16, respectively, indicating improved model fit (supplementary information Table S1). Adding an interaction term for social focus significantly improved the model fit compared to the null model for well-being ratings (*Chi*^2^ = 15.70, *p* = .015, AIC reduction = 4), but not for social connectedness and likeability ratings (Table S1). Adding an interaction term for personal focus significantly improved the model fit compared to the null model for likeability ratings (*Chi*^2^ = 14.87, *p* = .021, AIC reduction = 3), but not for social connectedness and well-being ratings (Table S1). These findings suggest that nationality has a broader and stronger influence on social and affective responses to movement synchrony, while personal and social focus have a more selective effect.

#### Social Connectedness

For social connectedness ratings, we found a main effect of the personal value universalism (*F*(1,341) = 5.44, *p* = .020), indicating that higher universalism was associated with higher ratings of social connectedness.

The value benevolence interacted with movement synchrony (*F*(2,341) = 7.06, *p* = .001; Figure 5A). Specifically, the self async / other sync slope was more negative than both the self sync / other sync (*z* = -3.36, *p* = .002) and the self sync / other async slopes (*z* = -2.44, *p* = .039), suggesting that compared to the other two movement synchrony conditions higher benevolence was more strongly associated with lower social connectedness ratings when the virtual-self moved asynchronously. Although the self sync / other async slope was more negative than the self sync / other sync slope, this difference was not significant (*z* = -2.23, *p* = .066).

**Figure 5:**
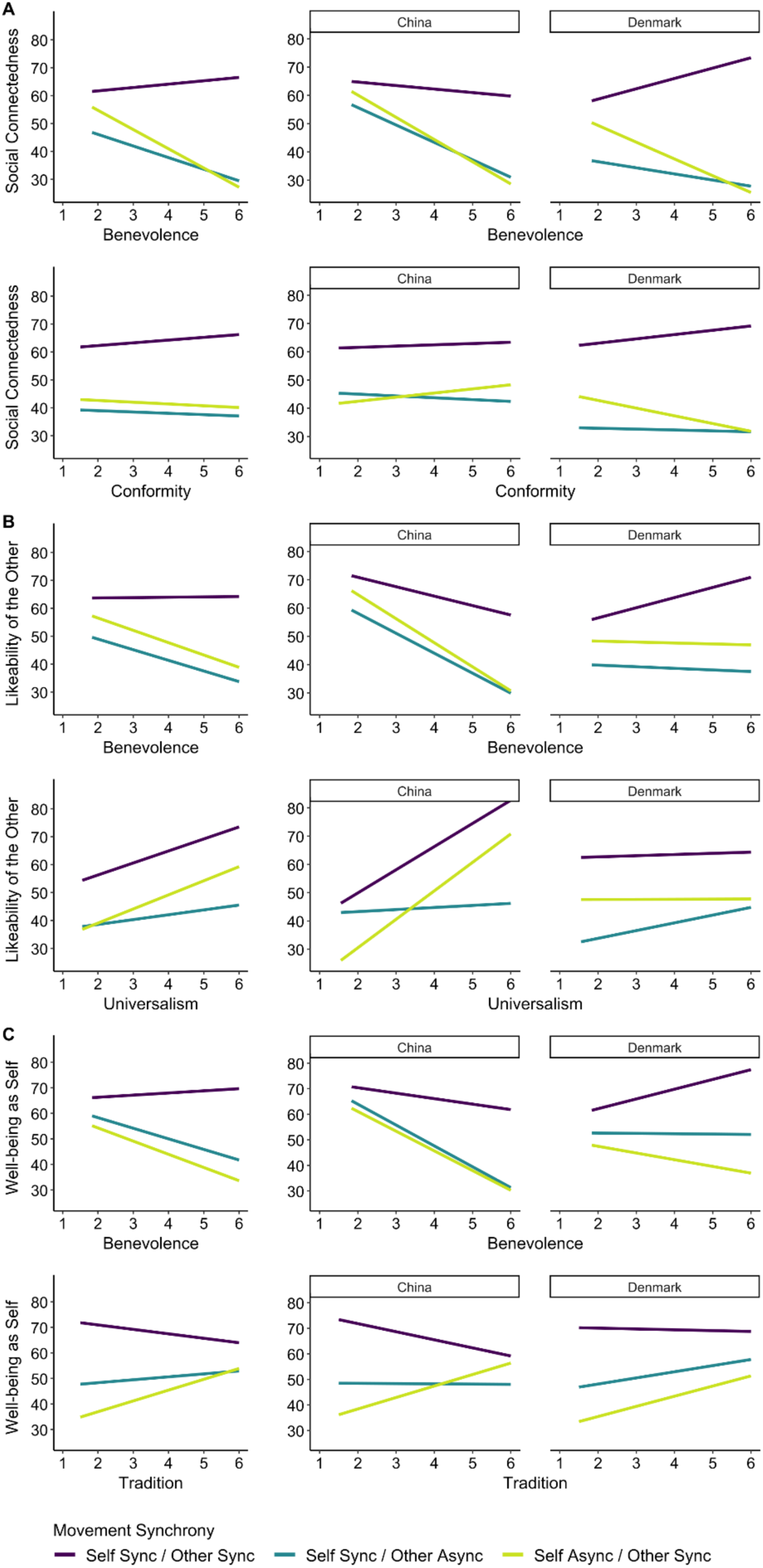
Model predictions from the exploratory analyses of personal values in the PVQ-RR. The displayed personal values are the ones with significant interactions with movement synchrony or three-way interactions with movement synchrony and nationality (*p* < .05). All other personal values showed no significant interactions.

The value conformity showed a three-way interaction with movement synchrony and nationality (*F*(2,341) = 3.08, *p* = .047; Figure 5A). The slopes of the self sync / other sync and self sync / other async conditions were slightly more negative for Chinese compared to Danish participants (estimates: -1.07 and -0.34), whereas the self async / other sync slope was more positive for Chinese compared to Danish participants (estimate: 4.19). This suggests that when the virtual self moved asynchronously, higher conformity was associated with higher social connectedness ratings in Chinese participants but with lower ratings in Danish participants. The pairwise comparisons of slopes between the two nationalities, however, were not significant.

#### Likeability of the Virtual Other

Likeability ratings showed an interaction between benevolence and movement synchrony (*F*(2,341) = 3.17, *p* = .043; Figure 5B), with the self sync / other sync slope being more positive than the self async / other sync slope (*z* = 2.47, *p* = .036), whereas the other two slope comparisons were nonsignificant (both *p* > .2).

Benevolence also interacted with nationality (*F*(1,341) = 5.76, *p* = .017; Figure 5B), indicating that Chinese participants who scored higher on benevolence rated the likeability of the virtual other lower, whereas Danish participants did not show such a trend.

The value universalism showed a three-way interaction with movement synchrony and nationality (*F*(2,341) = 3.17, *p* = .043; Figure 5B). The self async / other sync slope was more positive than the self sync / other async slope in Chinese participants (*z* = 2.60, *p* = .025) but not in Danish participants (*z* = -0.85, *p* = .674). All other pairwise comparisons were nonsignificant.

#### Well-being as the Virtual Self

For well-being ratings, the value benevolence interacted with movement synchrony (*F*(2,341) = 3.95, *p* = .020; Figure 5C). The self sync / other sync slope was more positive than the self sync / other async slope (*z* = 2.55, *p* = .029) and the self async / other sync slope (*z* = 2.44, *p* = .039), whereas there was no difference between self sync / other async and self async / other sync slopes (*z* = 0.49, *p* = .877).

Benevolence also interacted with nationality (*F*(1,341) = 5.01, *p* = .026; Figure 5C). Chinese participants who scored higher on benevolence rated their well-being lower, whereas Danish participants did not show such a trend.

Finally, the value tradition interacted with movement synchrony (*F*(2,341) = 3.90, *p* = .021; Figure 5C). The self async / other sync slope was more positive than the self sync / other sync slope (*z* = 2.79, *p* = .015), while the other two slope comparisons were nonsignificant (both *p* > .2).

## Discussion

This study examined the effects of interpersonal movement synchronization and music on social connectedness, likeability of a virtual other, and well-being of the virtual self in participants from individualistic (Danish) and collectivistic (Chinese) cultures. Using a video paradigm with virtual representations of self and other (Stupacher et al., 2017), we found that music, compared to a metronome, enhanced prosocial outcomes in both cultures, supporting the idea that music serves a fundamental social function across diverse cultures (Brown & Jordania, 2013). Consistent with previous studies (Stupacher et al., 2017; 2020; 2022), synchronized movements led to higher ratings of social connectedness, likeability of the virtual-other, and well-being of the virtual-self compared to asynchronized movements. While prior research has predominantly focused on Western, individualistic cultures, our replication of these effects in both Danish and Chinese participants suggests that the prosocial effects of movement synchrony are universal. However, cultural differences emerged in how participants responded to asynchronous conditions, highlighting the nuanced interplay between universal and cultural influences on different aspects of social bonding.

### Cultural Differences in Responses to Asynchrony

In ratings of social connectedness and likeability of the virtual-other, Chinese participants were more lenient toward asynchronized virtual-others, showing less negative evaluations compared to Danish participants. This aligns with our first hypothesis that collectivistic cultures prioritize social harmony and are more forgiving of misalignments to maintain group cohesion (e.g., Markus & Kitayama, 1991). For example, studies on communication in collectivistic cultures, such as Japan, have shown a greater tolerance for ambiguous social cues to preserve harmony (Gudykunst & Nishida, 2001). In China, the concept of *guanxi* emphasizes maintaining positive relationships and avoiding conflict (Leung & Wong, 2001), which may explain why Chinese participants rated asynchronous others less negatively. It is important to note, however, that the opposite outcome could have been equally well theoretically justified. As such, if an asynchronous other disrupts harmony, collectivistic individuals might evaluate such behavior more negatively, reflecting a different emphasis on maintaining social cohesion. Given the observed pattern of reduced negativity however, we interpret this as a result of conflict avoidance. Within collectivistic cultures, conflict avoidance is a well-documented strategy to maintain relational harmony and avoid overt criticism (Ohbuchi & Takahashi 1994; Bond & Smith 1996; Ting-Toomey & Kurogi 1998). From this perspective, less negative evaluations may reflect a deliberate down-regulation of criticism in order to preserve harmony.

Contrary to our second hypothesis, Chinese participants also rated their own well-being higher than Danish participants when the virtual-self moved asynchronously. While this initially appears inconsistent with the collectivist emphasis on social harmony, two complementary factors may help to explain this surprising pattern. The first factor is the difference in cultural inclinations to regulate affective responses towards norm deviations. In collectivistic societies, the concept of *mianzi*, (saving/preserving face) represents a central concern of the individual and promotes the regulation of self-presentation to maintain social harmony (Ho, 1976). This often results in down-regulation or suppression of explicitly negative self-evaluations in social situations. From this perspective, our finding of higher well-being as asynchronously moving virtual-self in Chinese compared to Danish participants does not necessarily indicate an absence of self-criticism, but may instead reflect a strategic regulation of emotional expression aimed at maintaining relational harmony.

A second, related factor concerns culturally normative response styles in self-reports. Cross-cultural research has shown how individuals from collectivistic cultures tend to report fewer negative self-directed emotions in self-report measures, even when internal self-evaluation may remain critical (Heine et al., 1999; Heine & Hamamura, 2007). The differences of well-being ratings in the self-asynchronous condition between Chinese and Danish participants may therefore reflect differences in how negative affect is regulated and expressed, rather than differences in the experience of norm violations per se.

In contrast, Danish participants exhibited stronger negative reactions to asynchrony, both for the virtual-other and self. This aligns with the individualistic emphasis on personal identity and competence, where asynchronous behavior may be perceived as a failure to meet personal or social standards (Heine et al., 1999). For instance, Yamagishi et al. (2008) found that individuals from individualistic cultures are more likely to attribute non-conforming behavior to personal dispositions rather than situational factors, which may explain the harsher evaluations of asynchronous others and selves among Danish participants.

### The Role of Personal Values

Because nationality and the PVQ-RR capture partially overlapping aspects of culture at different levels, their joint inclusion allowed us to assess which construct more strongly accounted for immediate social–evaluative responses. *Nationality* was a better predictor of social connectedness, likeability, and well-being ratings than the broad values of *social focus* and *personal focus* derived from the PVQ-RR. Importantly, while this suggests that cultural background encompasses additional, more relevant factors influencing different aspects of social bonding, it does not render personal individual values irrelevant. Rather, this distinction highlights the differences in how individual-level and cultural-level factors operate. While nationality indexes culturally internalized norms shaped through long-term socialization, the PVQ-RR primarily captures reflective and cross-situational value orientations, which may be too abstract to be directly relevant for the context-dependent social situation presented in the current paradigm.

At the same time, exploratory analyses of the 10 fine-grained basic PVQ-RR values provided further insights into how personal values may interact with cultural context. This supports the notion that personal values shape responses to movement synchrony, but also indicates that the aggregate PVQ-RR values may be too broad to capture the underlying effects. In this sense, specific PVQ-RR values reveal more nuanced modulations within cultural backgrounds.

*Benevolence*, which reflects caring for the welfare of close others (Schwartz et al., 2012), interacted with *movement synchrony* in all three ratings, and interacted with *nationality* in ratings of likeability and well-being. Higher *benevolence* was associated with lower social connectedness when the virtual-self moved asynchronously, suggesting that individuals who prioritize ingroup welfare may feel more distressed when they disrupt social harmony. Interestingly, higher *benevolence* was associated with lower likeability and well-being ratings in Chinese participants during asynchronous interactions but not in Danish participants. This discrepancy may reflect the conflict between Chinese participants’ social obligations and the stress of failing to meet them (Markus & Kitayama, 1991). In contrast, Danish participants, who may have more flexible social expectations (Oyserman et al., 2002), did not exhibit the same pattern.

*Conformity*, which involves avoiding upsetting others and complying with expectations (Schwartz et al., 2012), showed different effects across cultures. When the virtual-self moved asynchronously, higher *conformity* was associated with lower social connectedness in Danish participants but slightly higher social connectedness in Chinese participants. This suggests that *conformity* plays a more critical role in individualistic cultures, where individuals who care less about upsetting others may experience higher social connectedness when they disrupt harmony.

*Universalism*, which includes tolerance and societal concern (Schwartz et al., 2012), had a stronger positive effect on Chinese participants’ likeability rating when the virtual-other was synchronized. This suggests that Chinese participants may place greater emphasis on the actions of others in maintaining social harmony, while Danish participants were less influenced by *universalism*, reflecting their more individualistic orientation.

Finally, *tradition*, which reflects maintaining cultural and religious norms, influenced well-being ratings across both cultures. Participants with higher *traditional* values reported more stable well-being across synchrony conditions, while those with lower *traditional* values showed greater variation. This implicates that *tradition* may foster resilience in socially challenging situations.

In general, our findings highlight value-based differences in responses to social interactions. Personal values like *benevolence*, *conformity*, *universalism*, and *tradition* are expressed differently across cultures, underscoring the complexity of culture as a construct. While values help identify cultural orientations, their manifestations are influenced by cultural context, reflecting a dynamic interplay between values and culture (Hofstede, 2001; Schwartz, 1999).

### Limitations of the study

While our study provides valuable insights, several limitations must be acknowledged. First, reliance on self-reported measures introduces potential biases, such as social desirability and cultural differences in response styles (Chen et al., 1995; Harzing, 2006; Heine et al., 2002; van de Mortel, 2005). Future studies could incorporate interviews or training to enhance cross-cultural data comparability.

Second, conducting online research introduces challenges in experimental control and technical consistency, which may have affected participant engagement (Reips, 2002). We incorporated attention checks and excluded inattentive participants. However, future studies could benefit from more controlled environments to mitigate these issues. Additionally, although participants were raised and currently residing in their respective countries, accurately assessing their real-life cross-cultural exposure is difficult in an online study, which may have posed as a limitation, as such exposure could influence personal values or social attitudes.

Third, the reliance on only two nationalities limits the extent to which broader cross-cultural generalizations can be made. A more thorough design would ideally include multiple cultural samples, which would allow not only stronger inferences about cultural dimensions, but also reduce the risk that observed differences are due to mere chance or idiosyncratic features of the chosen samples (Schimmelpfennig et al., 2024). Future studies should therefore aim to incorporate broader cross-cultural designs to strengthen the validity of the cultural interpretations presented here.

Fourth, even though our findings are limited to Danish and Chinese participants, they represent specific cultural orientations. Cultural dimensions are multifaceted, and individualism and collectivism can be further differentiated into horizontal and vertical orientations (Singelis et al., 1995; Triandis & Gelfand, 1998). For example, Denmark’s horizontal individualism, which emphasizes autonomy and social equality, may differ from the vertically individualism of the United States (Nelson & Shavitt, 2002). Similarly, China’s vertical collectivism, which values hierarchy alongside group cohesion, may differ from the horizontal collectivism of other cultures. Future studies should include a wider range of cultural orientations to capture these nuances.

Fifth, the use of stick figures represents a trade-off between ecological and internal validity. While the minimalistic design reduced biases related to physical attributes, such as gender, age or ethnicity, enhancing internal validity (Blascovich et al., 2002), it was devoid of any contextual, relational or expressive cues that are present in real-world social interactions, reducing ecological validity and increasing ambiguity in participants’ interpretations. Given the consistent effects of movement synchrony across cultural groups and the comparable rates of failed attention checks between them, we do not interpret this reduced relatability as a confound that could lead to indifference or disengagement from the task. Rather, the abstract nature of the stimuli may have encouraged participants to rely more strongly on internalized social norms when evaluating concepts such as coordination and misalignment. From this perspective, future studies could further improve ecological validity by employing face-to-face interactions or virtual reality in multiple culturally diverse samples, thereby reducing interpretive ambiguity (Pan & Hamilton, 2018).

Finally, it is worth noting that although musical training did not significantly improve the statistical models, it differed between the two cultural groups. This difference may have influenced how rhythmic asynchrony was interpreted. The more musically trained participants in our sample may have been more accustomed to high rhythmic complexity and may therefore be more tolerant of asynchrony (e.g., Hannon & Trainor, 2007). Another possibility is that more musically trained participants may frame asynchrony as performance errors rather than a social irregularity. Future work could more directly assess how asynchrony is conceptualized across cultures or experimentally frame asynchrony as either a social or performance-related deviation.

## Conclusion

Our findings reveal both universal and culture-specific influences of movement synchronization and music on measures of social bonding. Synchronized movements and music universally enhanced social connectedness, likeability of a virtual other, and well-being, while cultural values and orientations shaped social dynamics to asynchrony. Chinese participants showed greater tolerance for movement asynchrony to preserve social harmony, whereas Danish participants exhibited stronger negative reactions to asynchrony, reflecting individualistic priorities. These insights highlight the potential of movement synchronization and music as universal tools for fostering social bonds, while underscoring the significance of cultural context in shaping interpersonal interactions. This has implications for enhancing cross-cultural communication and developing culturally sensitive therapies that leverage movement and music to promote social bonding and well-being.

## Data availability

The stimuli, data, and analysis scripts that support the findings of this study are openly available at https://researchbox.org/3886&PEER_REVIEW_passcode=LVBGRI.

## Ethical approval

The study was approved by the Institutional Review Board at the *** (****)

## Informed consent

Informed consent was obtained from all participants included in the study, and data protection, confidentiality and privacy were guaranteed.

## Author contributions

**: Conceptualization, Methodology, Investigation, Formal Analysis, Writing – Original Draft.

**: Investigation, Writing – Review & Editing.

**: Methodology, Writing – Review & Editing.

**: Methodology, Supervision, Writing – Review & Editing.

**: Conceptualization, Methodology, Investigation, Formal Analysis, Supervision, Writing – Original Draft, Writing – Review & Editing.

## Statements and declarations

### Ethical considerations

The study was approved by the Institutional Review Board at the Danish Neuroscience Centre (DNC-IRB-2023-010).

### Consent to participate

All participants provided written informed consent.

### Consent for publication

Not applicable.

### Data availability statement

The data that support the findings of this study, analysis scripts, and stimuli are openly available at https://researchbox.org/3886&PEER_REVIEW_passcode=LVBGRI.

