## Supplementary Information for "Cultural echoes in social bonding: Universal and culture-specific effects of movement synchrony and music on connectedness, likeability, and well-being"

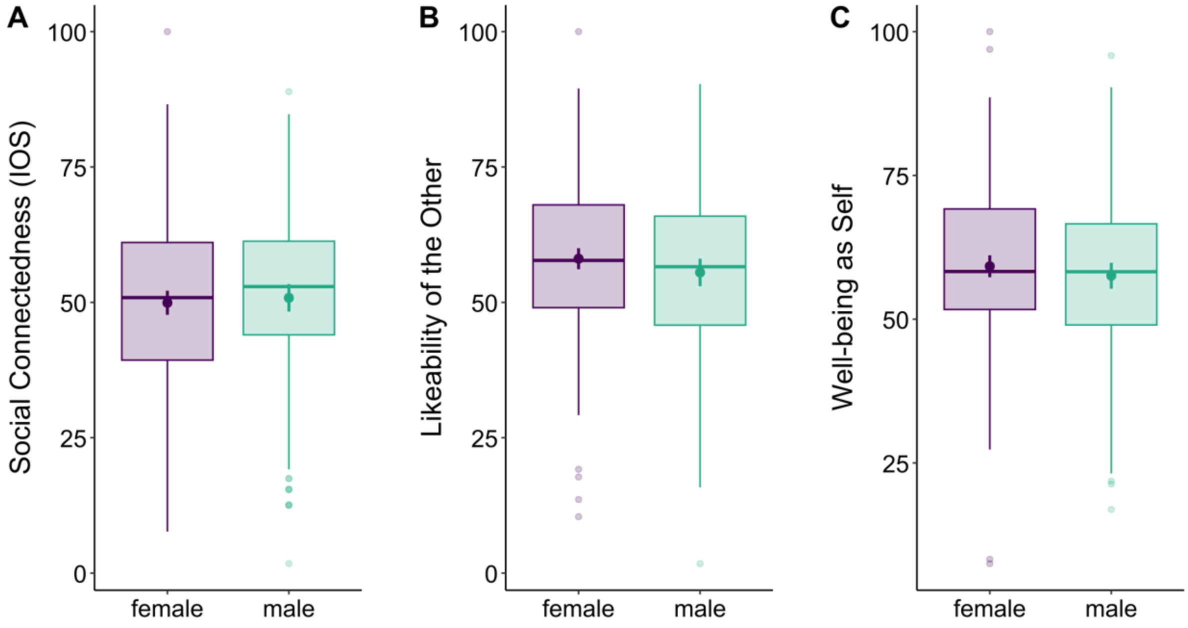


**Figure S1.** Mean ratings (●) with 95% confidence intervals (error bars) of social connectedness (A), likeability of the virtual other (B), and well-being as virtual self (C) for female and male participants. Two participants who identified as “other” were excluded from the plots. Boxes of the boxplots display the lower quartile, median, and upper quartile. Whiskers of the boxplots display data within 1.5 times the inter-quartile-range below the lower quartile and above the upper quartile. Dots represent datapoints outside this range.

**Table S1.** Comparisons of the null model *rating variable ~ synchrony * audio + empathy + bmrqSR + (synchrony * audio|participant ID) + (1|block)* with three models including either interactions between *synchrony * audio * nationality, synchrony * audio * social focus,* or *synchrony * audio * personal focus*. *** $p < .001$, ** $p < .01$, * $p < 0.05$. AIC: Akaike Information Criterion.

| Model | AIC | Comparison with null model | |
|---|---|---|---|
| | | $\chi^2$ (df) | *p* |
| IOS ~ synchrony * audio + empathy + bmrqSR (null model) | 37346 | — | — |
| IOS ~ synchrony * audio *** nationality** + empathy + bmrqSR | 37279 | 78.86 (6) | < .001*** |
| IOS ~ synchrony * audio *** social focus** + empathy + bmrqSR | 37346 | 12.26 (6) | .056 |
| IOS ~ synchrony * audio *** personal focus** + empathy + bmrqSR | 37349 | 9.36 (6) | .154 |
| Likeability ~ synchrony * audio + empathy + bmrqSR (null model) | 37925 | — | — |
| Likeability ~ synchrony * audio *** nationality** + empathy + bmrqSR | 37915 | 22.23 (6) | .001** |
| Likeability ~ synchrony * audio *** social focus** + empathy + bmrqSR | 37927 | 10.40 (6) | .109 |
| Likeability ~ synchrony * audio *** personal focus** + empathy + bmrqSR | 37922 | 14.87 (6) | .021* |
| Well-being ~ synchrony * audio + empathy + bmrqSR (null model) | 38440 | — | — |
| Well-being ~ synchrony * audio *** nationality** + empathy + bmrqSR | 38424 | 27.81 (6) | < .001*** |
| Well-being ~ synchrony * audio *** social focus** + empathy + bmrqSR | 38436 | 15.70 (6) | .015* |
| Well-being ~ synchrony * audio *** personal focus** + empathy + bmrqSR | 38447 | 5.20 (6) | .519 |
